# A side-by-side comparison of variant function measurements using deep mutational scanning and base editing

**DOI:** 10.1101/2024.06.30.601444

**Authors:** Ivan Sokirniy, Haider Inam, Marta Tomaszkiewicz, Joshua Reynolds, David McCandlish, Justin Pritchard

## Abstract

Variant annotation is a crucial objective in mammalian functional genomics. Deep Mutational Scanning (DMS) is a well-established method for annotating human gene variants, but CRISPR base editing (BE) is emerging as an alternative. However, questions remain about how well high-throughput base editing measurements can annotate variant function and the extent of downstream experimental validation required. This study presents the first direct comparison of DMS and BE in the same lab and cell line. Results indicate that focusing on the most likely edits and highest efficiency sgRNAs enhances the agreement between a “gold standard” DMS dataset and a BE screen. A simple filter for sgRNAs making single edits in their window could sufficiently annotate a large proportion of variants directly from sgRNA sequencing of large pools. When multi-edit guides are unavoidable, directly measuring the variants created in the pool, rather than sgRNA abundance, can recover high-quality variant annotation measurements in multiplexed pools. Taken together, our data show a surprising degree of correlation between base editor data and gold standard deep mutational scanning.

## INTRODUCTION

A major goal of mammalian functional genomics is to understand the genotype-phenotype relationship (1). However, achieving this lofty goal requires large-scale experiments that can, in parallel, examine both genotypes and phenotypes in pooled assays. While gene level loss-of-function screens in mammalian cells rose to prominence with the invention of RNAi based screening tools (2–4) in the early 2000’s and eventually CRISPR/CRISPRi (5–8) in 2012-2013, the high throughput annotation of individual coding variants in mammalian cell lines is a more recent innovation (9, 10).

A leading technology in the field of high throughput variant annotation is a technique known as deep mutational scanning (DMS) (10). Deep mutational scanning libraries tend to involve large cDNA libraries of single amino acid mutations that encompass all 20 possible amino acids at every position.

When performed in mammalian cell lines, these libraries are introduced via a transduction (11–13), or into a safe harbor “landing pad”(14–16). While DMS is capable of providing comprehensive measurements of variant effects (10, 17, 18), DMS has nonetheless been difficult to scale to large genes or to multi-gene families, and the measurements may not reflect the effects of the same mutations at the endogenous genomic locus. Moreover, the technical challenges involved in DMS can lead to variable dataset quality (19).

As a functional genomic technology, base editing is in its early stages. Base editing (BE) screens use nCas9 to target a deaminase to a specific site in the genome and generate transition mutations (C>T or A>G) (20, 21). BE screens use a surrogate measure of genotype by sequencing the guide RNA (gRNA) sequence, which allows BE screens to measure phenotypes across the genome (22–24). Moreover, base editor screens have other major advantages that can include the ability to edit at the endogenous genomic locus and the ability to identify splicing defects (22, 25–28). However, base editor screens also present certain challenges. The primary challenges are: 1. BE efficiency (only a portion of individual cells harboring an sgRNA are likely to be edited and some cell lines do not edit well (29)), 2. off-target editing (while editing is largely constrained to a small window within the sgRNA non-target sequence, some off-target editing is likely occurring (30, 31)), 3. bystander editing (when more than one possible edit occurs in an on-target editing window, the amino acid variant(s) made are more challenging to infer (32, 33)), and 4. protospacer adjacent motif (PAM) requirements (PAMs limit where sgRNAs targets (34) and PAM- less variants appear to have decreased efficiency (35–37)). These issues have led the field to view BE screens as a method for initial identification of interesting variants and regions but with limited capability to directly annotate loss-of-function phenotypes.

When competing high throughput measurement methodologies can generate similar data, it can be extremely useful to benchmark these methods against each other. For instance, the direct comparison of CRISPR Cas9 LOF screens and RNAi screens suggested that CRISPR is a more sensitive and specific technique for identifying essential LOF phenotypes, but that RNAi screens can help understand the dosage sensitivities of essential genes and can sometimes rescue false negatives in CRISPR screens for a subset of biological functions (38–40). Additionally, the direct comparisons of high throughput drug sensitivity measurements found that the precise metrics and methods that are used in comparing datasets can create different conclusions on dataset reliability and usability. Together, these high-profile efforts highlight the importance of a careful comparison of high throughput datasets using multiple metrics and the public dissemination of the resultant data (41, 42).

Here we perform the first direct comparison between base editing and deep mutational scanning in the same cell line in the same lab. To accomplish this, we use the Ba/F3 cell system. This allows for a direct comparison and eliminates differences in genetic context as a confounding variable driving the differences in measurements between the approaches. Using this system, we identify specific data filters that generate largely matching conclusions about the phenotypes of loss-of-function variants. We also identify a two-step high throughput workflow for base editor screens that can streamline the validation of variant interpretation in pools by directly sequencing the edited variant fraction with error corrected sequencing (43).

## MATERIALS AND METHODS

### DMS Library Preparation and Screen

BCR-ABL cDNA was cloned downstream of EGFP in the pUltra (Addgene #24129) lentiviral vector by GenScript to make pUltra BCR-ABL WT (Addgene #210432). Twist Bioscience generated a saturating mutagenesis (SM) library of single amino acid changes in the N-lobe of the ABL kinase domain. NEB Stable chemically competent (NEB #C3040I) cells were transformed with the SM library, with a coverage of >1000X, on to 15-cm LB agar plates with ampicillin. After 48 hours at 30°C, colonies were scrapped off the agar and plasmid DNA was extracted using Omega Bio-Tek’s E.Z.N.A. Plasmid DNA Midi kit.

HEK293Ts were transfected with 35 ug of SM ABL library and 10 ug of helper plasmids (1:1:1) in 10-cm dishes using Thermo Fisher Lipofectamine 3000 (5 Lipo : 1 DNA). The next day the media was changed to fresh RPMI (Cytivia SH30027.02). After 36 hours viral RPMI media was used to infect BaF3s, in the presence of 1 ug/mL mouse IL-3 (peprotech 213-13) and 6 ug/mL polybrene, at a low multiplicity of infection. After another 36 hours, BaF3s were maintained in RMPI with 1 ug/mL IL-3. Infected cells were enriched by fluorescence-activated cell sorting on EGFP at the Penn State Flow Cytometry Core facility.

At the start of the DMS Screen, IL-3 was removed, and 30 million cells were saved to establish a baseline mutation frequency. Approximately 5 million cells were treated with DMSO for 6 days. Media was refreshed on day 3. Cell count was tracked by a BD Accuri C6 Plus flow cytometer. Cells were maintained in exponential phase. If cell viability was less than 90%, then viable cells were enriched by Ficol-Paque (Cytiva).

### DMS Library preparation and Single Strand Consensus Sequencing

High quality genomic DNA was extracted by Monarch Genomic DNA Purification Kit. Then a modified and scaled-up CRISPR-DS workflow was used to determine accurate variant distributions (43). For each time point 20 ug of genomic DNA specifically was digested just outside of the mutagenized region of ABL kinase cDNA using Cas9. After end-repair and A-tailing, UMI ligation was used to label single molecules of DNA. Then NGS indices were added by PCR to help deconvolute samples after pooling. Oligo enrichment and PCR were used to further increase the abundance of the mutagenized region in the pooled samples.

One-hundred-fifty-nucleotide paired-end sequencing of the UMIs and mutagenized region was done on an Illumina NovaSeq 6000. For each sample, dunovo was used to generate error-corrected single strand consensus from the UMI barcodes. Then bwa-mem2 was used to align the census to human ABL cDNA. After filtering out for mouse ABL reads, aligned reads with less than 5 mismatches would undergo variant calling and annotation using a custom R script. Briefly, for each alignment, variants were converted from the MDZ read tag and normalized to read depth at that position. Mutant growth rates were calculated using exponential growth equation and the mutant allele frequency (MAF):

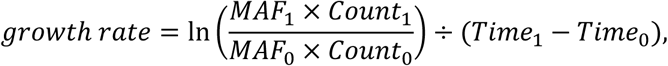

where the subscripts 0 and 1 denote the initial and final time point, respectively. Cell counts and the splitting ratio were used to account for dilution due to cell splitting during the DMS screen. Time is measured in units of hours. Skewed Gaussian mixture models were fit over the bimodal distribution of mutant growth rates using the Curve_fit function from the Scipy python package (44). Z-score cutoff for DMS data was determined by fit mean and standard deviation of the wild type like component of the mixture distribution.

### Base Editor Library Preparation, Screen, and Sequencing

BCR-ABL tiling guide RNA (gRNA) sequences were generated by CHOP-CHOP (45) with ‘NGN’ PAM setting. Guides were cloned into lenti-sgRNA hygro vector (Addgene #104991) by GenScript to make the BCR-ABL sgRNA library. ABE8e SpG plasmid was made by deleting the U6 sgRNA cassette from pRDA_479 (Addgene #179099) (46) using NEB KLD (NEB #M0554S).

Ba/F3s infected with pUltra BCR-ABL WT (Addgene #210432) were allowed to grow in the absence of IL-3. After 5 days >95% of the BaF3s were EGFP+. Then they were infected with either ABE8e SpG or CBEd SpG and selected with 1 ug/mL puromycin for 5 days. In a 15-cm dish HEK293Ts were transfected with 60 ug of the BCR-ABL and control sgRNA libraries and 40 ug helper plasmids (1:1:1:1) using calcium phosphate. The next day DMEM media was changed to RPMI. Independent infections of pUltra BCR-ABL WT ABE8e BaF3s were done at low multiplicity of infection and >500X coverage.

Transfected HEK23Ts were saved to establish a baseline sgRNA frequency. Three days after sgRNA library infection, the cells were selected with 1 mg/mL hygromycin for 6 days and pelleted. Genomic DNA was extracted using phenol-chloroform (47) and quantified by Qubit. Staggered PCR was used to extract sgRNA sequences from 10 ug of genomic DNA, as described previously (48) with two modifications: custom forward staggered primers were used (Supplemental Table 1) and PCR amplicons were gel extracted using Omega Bio-Tek Gel Extraction kit.

### sgRNA Analysis

Guide RNA amplicons were trimmed with CutAdapt (49) to remove the U6 promoter and gRNA scaffold. The remaining sequences were aligned to a reference list of guides using Bowtie (50). Guide counts were established by Counter from the Collections python package. pyDESeq2 (51) was used to determine the fold change in guides between time points. The fold change was converted to growth rates using the formula above assuming wild-type growth rate. Z-score cutoff for ABE data was based on the human targeting (AAVS1, CCR5, and ROSA26) control sgRNA.

### Verification Screen, Sequencing, and Analysis

A pool of 20 guides were synthesized by IDT and cloned into lenti-sgRNA hygro vector using golden gate sites (48, 52). After making BCR-ABL WT ABE8e BaF3 cells as above, IL-3 was returned to the RPMI media. The lentivirus library of sgRNAs was made by calcium phosphate transfection of HEK293T. Two independent infections of 1.5 million BCR-ABL WT ABE8e BaF3 cells grew for 3 days before the start of 1mg/mL hygromycin selection. After 6 days, hygromycin was removed, a pellet was saved for sgRNA sequencing, and the cells recovered for 2 days. After the 2-day recovery, IL-3 was removed, and a pellet was saved for mutation sequencing. Six days after IL-3 withdrawal pellets were saved for mutation sequencing. Nineteen days after IL-3 removal pellets were saved for sgRNA sequencing. Mutation and sgRNA sequencing were performed as described above.

A custom Python script was used to determine the frequency of in phase or *cis* mutations made by ABE. Briefly, UMI_tools (53) was used to move 24 nucleotide UMI in to read header. At least 5 million paired end reads were aligned to the ABL kinase after 12 nucleotides were hard clipped from the 5’ end using Bowtie2 (54). To deduplicate barcodes, each sample’s barcodes was grouped based on alignment with an allowance of one mismatch using Bowtie (50). Only the most frequent UMI in each group continued to the rest of the analysis. Mismatches with a quality score greater than 30 were grouped by UMI. A mutation was called and counted if the mismatch is observed in more than 80% of reads in the UMI, and the UMI contains more than 2 reads. Fold change was determined by pyDESeq2 (51) and converted to growth rate using WT growth rate. Background mutation frequency was set based the 95^th^ percentile of non-(A > G) and non-(T > C) mutant frequency outside of the 2 to 12 nucleotide sgRNA editing window. All A > G mutations within the targeting strand and editing window of an sgRNA were linked to that sgRNA.

### BE-HIVE Weighted Model

In order to determine how well the on-target editing sgRNA phenotype can be recapitulated, sgRNA and editing efficiency was predicted by BE-HIVE (55). Only edits with a frequency greater than 0.05 and sgRNAs with completely matched DMS measurement were allowed. The growth rate of an sgRNA is sum of the relative growth rates of edited and un-edited cells:

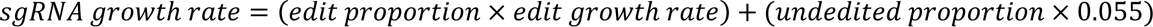

In the case of edits across multiple amino acids, a null model of mutant interactions predicts the growth rate of the multi amino acid mutant as the product of individual mutants divided by the *WT* growth rate multiplied by the *WT* growth rate (0.055 hour^-1^) (56). In other words, two deleterious *cis* mutations are expected to be more deleterious than the individual mutants. If there was an epistatic interaction, the growth rates of the double mutant would differ from the null model.

## RESULTS AND DISCUSSION

### DMS Data is high quality and correlates with Evolutionary Conservation, Secondary Structure, and Function

For our comparison experiments, we chose to use a fit-for purpose cellular system, where the growth of the cell line, Ba/F3, depends upon the presence of IL-3 in the media until an activated tyrosine kinase is added. This system allows us to introduce the DMS library in the same genomic context as the target of the base editor screen. Heterologous expression of a tyrosine kinase cDNA integrated into the genome is advantageous with respect to our study design because all variants (DMS or BE) are evaluated using the same promoter. We use the BCR-ABL oncogene as the activated tyrosine kinase for several reasons: 1. It is an important oncogene that is of interest to applied and basic research groups. 2. Its structure has been solved many times in many conformations (57, 58). 3. There are years of experiments that can be drawn from to gain confidence in the resultant data (58–60).

To perform the deep mutational scan (DMS) in BCR-ABL, we designed a saturation mutagenesis library spanning amino acid residues 242 to 320 in the N-lobe of the ABL1 kinase domain. This region was chosen for its reasonable size and the presence of known structural features that include the p-loop, the gatekeeper, and the alpha-C helix. This library contained 1441 AA variants and was transduced into Ba/F3s in the presence of IL-3. After infection, fluorescence activated cell sorting (FACS) was used to enrich for infected cells. Following recovery from sorting, the Ba/F3 cells were screened for six days without IL3. In this negative selection screen, variants expressing non-functional copies of the BCR-ABL fusion protein deplete from the population. After the screen, we perform a sensitive barcoded sequencing protocol where the heterologous copy is specifically excised from the genome using Cas9- sgRNA complexes that are specific to the cDNA (43). These genomic fragments are then ligated with unique molecular identifiers to allow for the deconvolution of library variants and the elimination of PCR and sequencing noise (Figure 1A). Initially, the library contained ninety seven percent of all possible single residue changes from AA 242-320.

**Figure 1.**
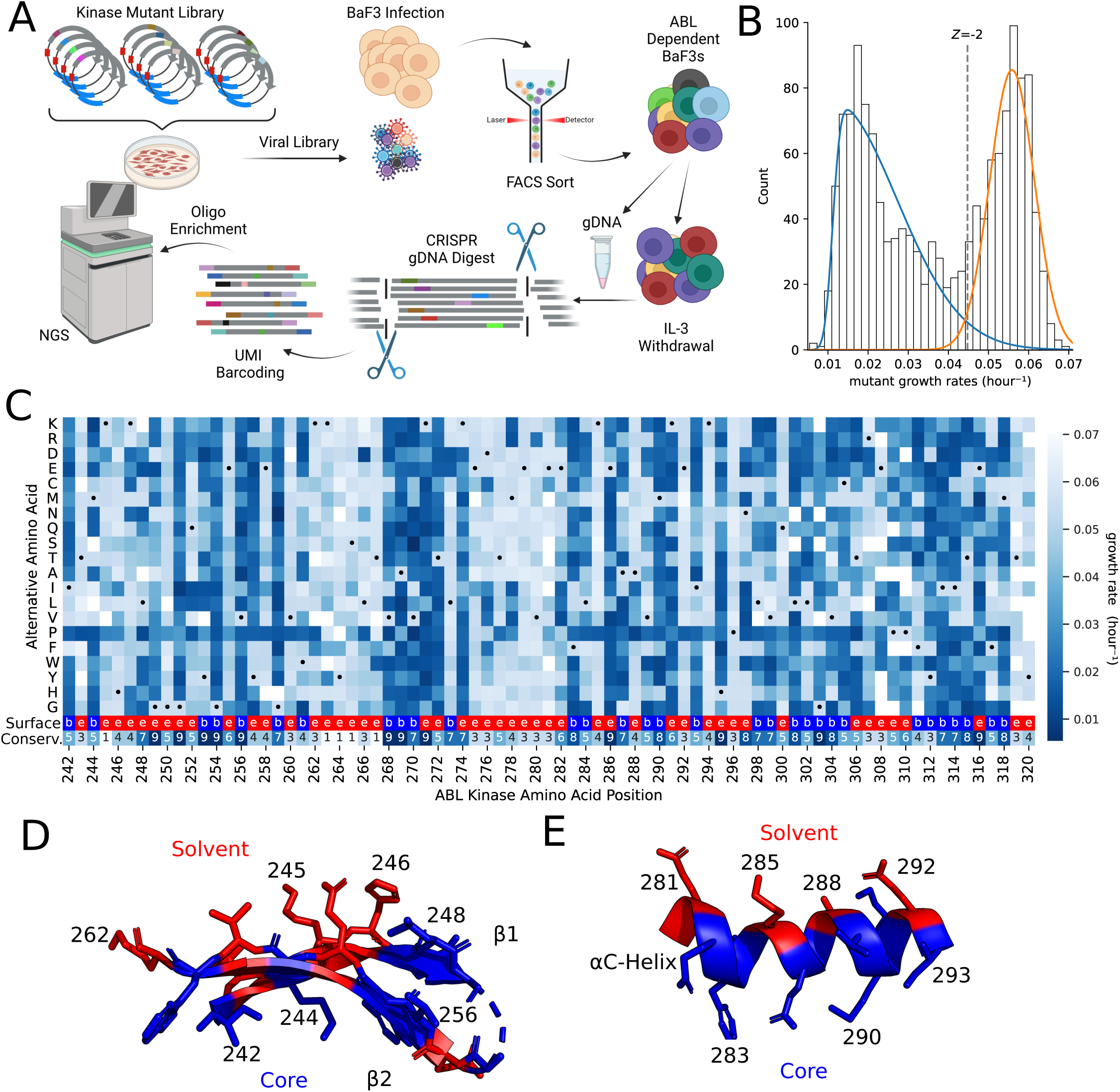
Functional landscape of ABL N-Lobe. (A) Schematic of deep mutational scanning. After lentiviral integration of EGFP-P2A-BCR-ABL, BaF3s were sorted to enrich EGFP+ infected cells. Cells were pelleted before and 6 days after IL-3 withdrawal. After targeted genomic DNA digest by Cas9, single molecules of DNA were labeled by UMI ligation. Then biotinylated oligo baits were used to enrich the mutagenized region. (B) The distribution of mutant growth rates in the ABL N-lobe is bimodal. Two skewed gaussians are fit to determine the variation in deleterious (blue) and “wild-type like” (orange) mutations. The dotted line represents a -2 Z-score threshold with respect to the “wild type-like” distribution. (C) Heatmap of the growth rate of mutations at each position in ABL1 N-lobe. Black dot represents WT positions. Missing data is in white. Second to the last row of the heatmap provides surface exposure information because solvent exposed residues (red, “e”) tend to be more tolerant of mutations than buried residues (blue, “b”). Tolerance/sensitivity to mutagenesis is projected onto two key structural features of the ABL N-lobe (PDB 6XR6): the (D) anti-parallel beta-sheet, and the (E) αC- Helix. If mean growth rate of alternative alleles at a residue is less than the -2 Z-score cutoff, then the residue is colored blue. In contrast, if the mean growth rate of alternative alleles is greater than the –2 Z-score cutoff, then it is colored in red.

When the mutant in BCR-ABL impairs kinase function, then that cell will exhibit a reduced growth rate. After sequencing, we found that mutant growth rates form a bimodal distribution of counts, consistent with measured distributions of fitness effects in other DMS screen (12, 15, 16). One of the peaks in this bimodal distribution of growth rates centered at approximately 0.02 hour^-1^ and constituted the set of deleterious mutations, while the other distribution mode was centered at the known *WT* growth rate of 0.055 hour^-1^ and harbored AAs that are “wild type like fitness” (Figure 1B). To call the “hits” of specific amino acid variants that are required for growth, we use a Z-score cutoff of -2 for the “wild type like” distribution. This specific cutoff is used to be consistent with prior literature in the sgRNA community (23, 24, 46, 61) and to draw a line that can be approximated in both studies using approximately analogous statistical and experimental criteria (Figure 1B). Using this cutoff, we estimate that 56% of measured variants in the N-lobe of the ABL kinase impair kinase function.

Initial observations of the deleterious residues in the DMS dataset are consistent with prior knowledge. For instance, the catalytic lysine K271 is required for kinase activity and mutations at that position strongly deplete (measured growth rate averaged 0.019 hour^-1^) (Figure 1C). Moreover, the systematic insertion of prolines during DMS studies causes a discernible “proline band” in high quality studies (12, 62). We clearly observe a proline band in our data (Figure 1C). Beyond these critical depletion signals, a known flexible and non-conserved region in ABL1 (i.e. residues 262-267 between the β2 and β3 strands) did not deplete.

More systematically, functionally important residues are typically evolutionarily conserved and their mutagenesis tends to be deleterious in DMS studies (10, 16). Consistent with this, there is a -0.74 correlation (p < 0.001) between the mean mutant growth rate at a residue and the evolutionary conservation scores of that residue from ConSurf (57, 63–65)(Figure 1C, bottom row of heatmap, Supplemental Figure 1). Moreover, from a biophysical perspective, solvent accessible residues tend to be more tolerant of mutagenesis and are less conserved (66). We observe this trend in Figure 1C, 1D and Supplementary Figure 1.

The ABL N-lobe is composed of conserved structural elements that include the alpha-helix called the αC-helix, the five-stranded antiparallel β-sheet, the GxGxxG motif of the P-loop, as well as several catalytically important residues. Investigation of these key structural features reveals patterns that are consistent with known structure-function relationships. The GxGxxG of the p-loop spans residues 249- 254 and all 3 conserved glycines show strong depletion phenotypes. Additionally, the alternating banding pattern in anti-parallel β -strands 1 and 2 (residues 242-262, Figure 1D) is explained by essential residues facing towards the substrate pocket (blue residues, Figure 1D). Moreover, the αC-helix, dynamically transitions from its “in” to its “out” state during kinase activation. This highly conserved regulatory dynamic requires support from the underlying hydrophobic core. Thus, the residues that interact with the core of the protein show the expected pattern of intolerance (blue, Figure 1E), while the outward facing residues show tolerance to mutations (red, Figure 1E). It is the connection to known structure, function, and conservation data that gives us high confidence in the validity of the DMS data as a gold standard for comparison with sgRNA data.

### An Adenosine Base Editing Screen Isolates Functionally Important Domains and Residues

One of the major benefits of base editor (BE) screens is the ability to rapidly screen across a protein’s entire length. To demonstrate this breadth and speed, we rapidly cloned a tiled library of 3115 sgRNAs across the entire BCR-ABL cDNA, including regions beyond the original DMS library. The BE phenotype was determined by the change in sgRNA abundance following exponential growth in cell culture (Figure 2A). Looking across the entire length of the protein (Figure 2B), we performed a sliding window estimate of the proportion of strongly negative Z-scores across the protein (see Methods). This sliding window estimate highlights the regions of the protein that are unusually enriched for the presence of highly essential residues. Colored regions corresponding to the coiled-coil (CC) domain, double homology (DH) domain, plekstrin homology (PH) domain, Src-homology 3 (SH3), Src-homology 2 (SH2), the kinase domain, and F-actin binding domain (FAB) are highlighted. Highly significant domains that have been previously implicated in transformation, such as the CC (67, 68), SH2 (68–70), Kinase (71), and FAB domains (72, 73) show significant depletion. While the role of the DH domain is not understood, our base editor data suggests that it is important for growth factor independence (Figure 2B) (74, 75). Finally to call depletion “hits” we used a –2 Z-score based on the distribution of negative-control guides to be consistent with Hannah et al. (23) (Figure 2C). Applying this criteria yields 387 hits, or approximately 12.4% of all BCR-ABL sgRNAs (consistent with others (23, 46)). Most notably, there is a strong over-representation of depletion phenotypes for guides that target the ABL kinase domain, with 38% of kinase domain guides depleting below a –2 Z-score. In concordance with the DMS screen above, the guides that can target the conserved regions of the P-loop, Lys 271, and the buried region of the αC- helix produce deleterious phenotypes. (Figure 2D).

**Figure 2.**
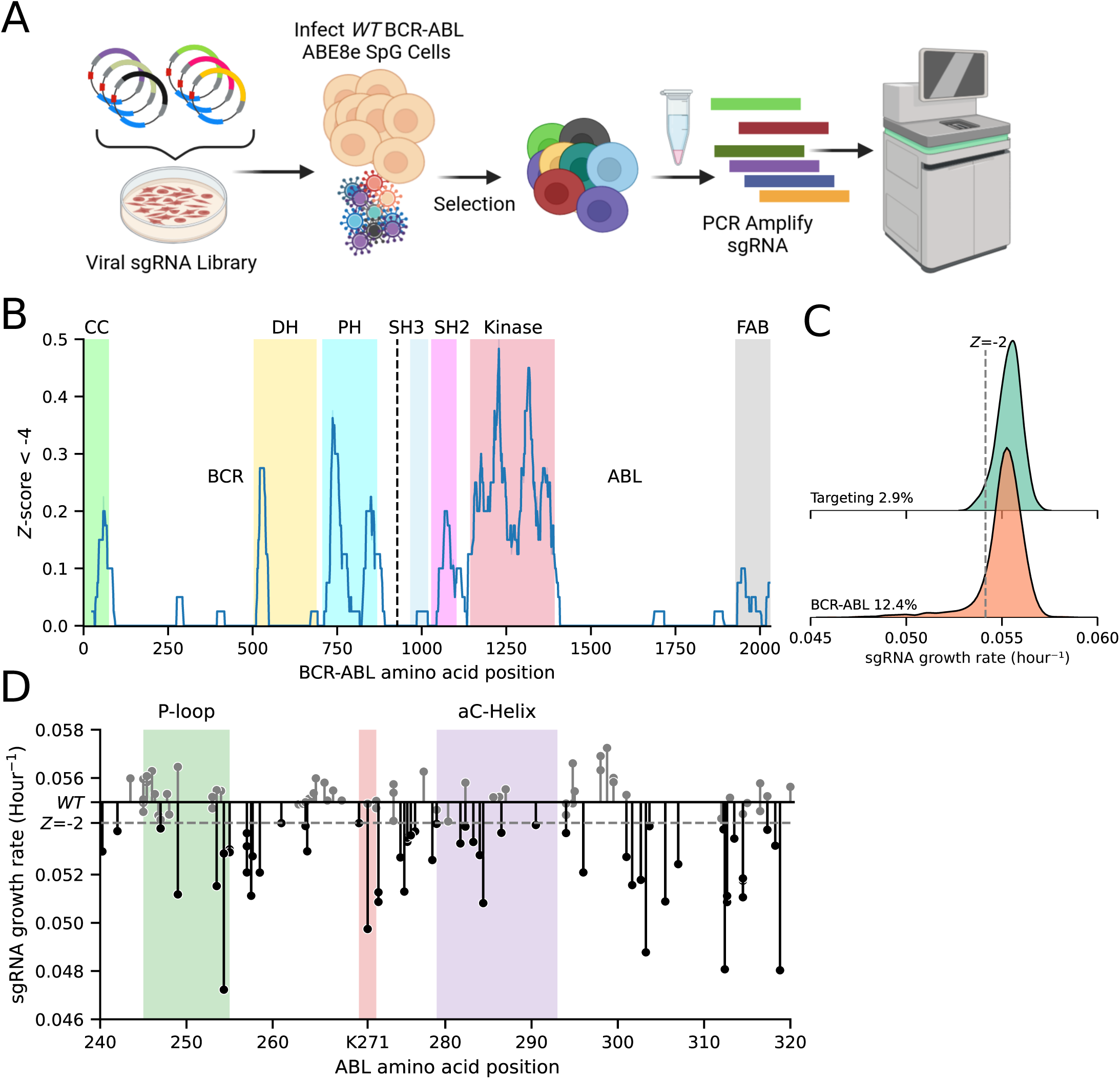
Adenosine Base Editor screen of full length of BCR-ABL. (A) Schematic of adenosine base editor screen. Three days after infection with sgRNA library, EGFP-P2A-BCR-ABL1 ABE8e BaF3 were selected with 1mg/mL hygromycin for 6 days and pelleted. Guides were PCR amplified and sequenced. (B) Sliding window of 40 sgRNAs estimate of the proportion of BCR-ABL sgRNAs that drop out more than a Z-score of -4 of the non-targeting control sgRNA growth rate. (C) Kernel density estimate of growth rate distributions of non-targeting control, and BCR-ABL sgRNA libraries. Dashed grey line represents a – 2 Z-score of the targeting control. (D) Lollipop plot displays dropout of each sgRNA across the ABL kinase domain. Dashed grey line represents a –2 Z-score of the targeting control.

### Side by side comparison of ABE predicted edits and DMS Screens

To assess the correlation between our ABE screen and deep mutational scan, we compared growth rates of sgRNAs and their predicted edits to growth rates of variants from the deep mutational scan. We focused on 80/118 sgRNAs for which all predicted edits have variant information in the DMS screen. (Figures 1C and 2D).

The predicted edits of the sgRNAs predominantly occur within an editing window (32, 76). While the editing window is in the non-targeting strand, we (and others) refer to the window relative to the guide sequence position. Most edits occur between positions 2 and 12 (76, 77). Plotting sgRNA data against DMS data (Figure 3A), each dot represents an individual variant, with each sgRNA appearing as a row of dots for all of its possible edits. Notably, the dynamic range of the Y-axis is reduced for base editing compared to DMS. (there is an ∼ 0.045-0.06 hour^-1^ Y-axis measurement range for base editing versus ∼0.01 to 0.06 hour^-1^ for DMS on the X-axis). This is likely because counting an sgRNA read can count either edited or unedited cells.

**Figure 3.**
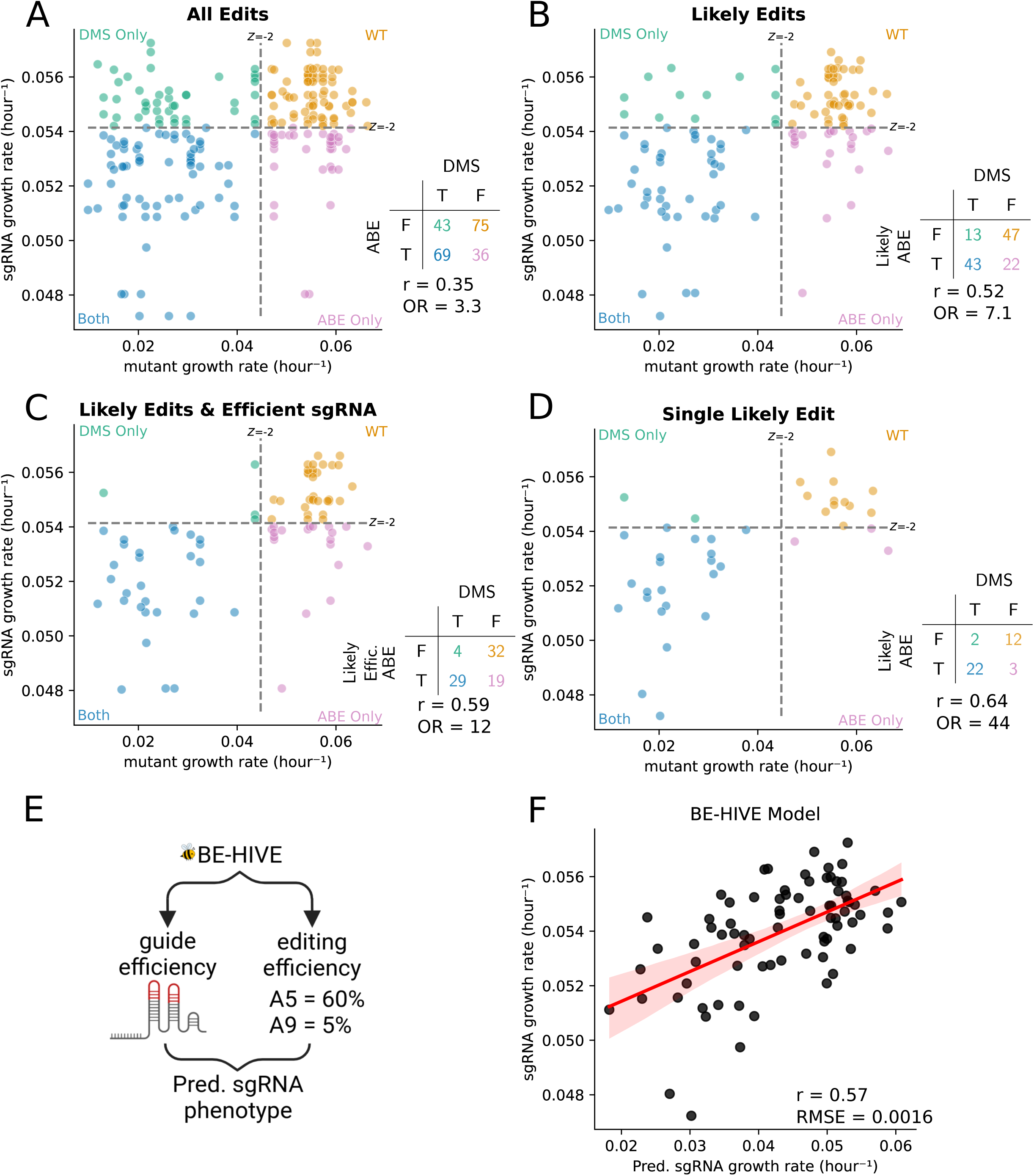
Comparison of Adenosine Base Editor sgRNA growth rate and their respective mutation growth rates from Deep Mutational Scan. Each dot represents a mutation an sgRNA is predicted to make. Dashed lines represent –2 Z-score of the non-deleterious distribution and non-targeting control sgRNA for the DMS and ABE screens, respectively. These cutoffs are used to define if an sgRNA or mutation is deleterious. If an sgRNA and its mutation does not deplete in their respective screen, in other words both are non-deleterious, then they are colored orange. If they both are deleterious, or true positive, then they are colored blue. If an sgRNA deletes, but the predicted edit does not deplete, a false positive, then the dot is colored in the pink. If a sgRNA fails to deplete and the predicted mutation(s) are deleterious, a false negative, then that point is colored in green. (A) shows all possible edits between nucleotides 2 and 12. (B) shows only the most likely edits, those between nucleotides 4 and 8. (C) shows only sgRNA predicted to be efficient and edit between nucleotides 4 and 8. (D) shows sgRNAs that are predicted to make only a single edit between nucleotides 4 and 8. (E) The distribution of edits can be estimated by machine learning model called BE-HIVE. (F) Correlation between predicted sgRNA growth rate and observed sgRNA growth rate. The x-axis shows the predicted growth rate of each sgRNA based on a weighted sum of the probability edit(s) and the effect of that edit(s) from DMS data. The y-axis shows the measured growth rate of the efficiently editing sgRNAs from the ABE screen.

This initial analysis yields a modest but significant Pearson correlation of 0.35 (Figure 3A). This indicates a relationship but suggests caution in directly annotating individual variants from a raw sgRNA experiment without further validation of the variants with individual sgRNAs.

Considering the modest correlation with all putative variants, we hypothesized that the 2-12bp editing window was too broad. By focusing on a narrower, more efficient 4-8 nucleotide editing window (23, 77, 78), the Pearson correlation improved from 0.35 to 0.52 and the odds ratio of hits versus non-hits increased to 7.1 (Figure 3B). This improved agreement came at the cost of removing 26 potentially correctly annotated deleterious variants (Figure 3A, 3B and Supplemental Figure 2).

After applying the “likely” edits filter to generate Figure 3B we still identify 13 false negative and 22 false positive variants in our plots. In terms of these remaining false negatives, an sgRNA could fail to deplete when sgRNA sequences are of low efficiency (48). Thus, we examined the consequences of adding an additional filter requiring an sgRNA efficiency score greater than 50 (Figure 3C, Supplemental Figure 2S) (48). This simple additional cutoff further improves the Pearson correlation between sgRNA predicted edit growth rates and DMS data to 0.59 and eliminates 9 of the 13 remaining false negative variants (Supplemental Figure 2S).

Next, focusing on the false positive variants in Figure 3B (pink dots), we identified 15 sgRNAs that are predicted to make 22 false positive variants. Interestingly, 10 of the 15 sgRNAs are multi-edit sgRNAs that are predicted to also make true positives edits (blue variant dots). One simple way to reduce the effect of multi-edit ambiguity is to only examine the sgRNAs that are predicted to make a single edit in the 4 to 8 nucleotide editing region. The addition of this “single likely edit” filter enhances the correlation to 0.64 (p-value < 0.001) and the OR for hits to 44 (p-value < 0.001) (Figure 3D). For single-likely-edit sgRNAs there is a true positive rate of 0.88 and an accuracy of 0.87 with respect to gold standard DMS data. However, this filter removed 14 out of 39 original sgRNA hits. For detailed information on filters and filtered sgRNAs/variants, see Supplementary Figures 2 and 3.

Given that filtering sgRNAs by efficiency, edit probability, and the number of edits improves annotation confidence at the expense of total hits, we sought to more effectively utilize data from multi-edit sgRNAs (Figure 3D). At a first pass, it seems like one might create variant-level interpretations for multi-edit sgRNA hits just by predicting the ensemble of sgRNA edits and their abundance in the population using machine learning algorithms like BE-HIVE (55). However, while BE-HIVE can predict the ensemble of mutations (Figure 3E), it can’t predict the proportion of the measured phenotype that is attributable to those mutations. Therefore, we investigated whether multi-edit sgRNA dynamics could be predicted from the combined dynamics of their polyclonal edits. We used the BE-HIVE predicted allele frequencies vector (to approximate the structure of the polyclonal population) and simply assumed that all of the predicted variants grow according to the gold standard DMS growth rates. This estimated the expected multi-edit sgRNA growth rate, providing insight into the contribution of in-window editing versus off-target effects to sgRNA dynamics.

We observed a strong correlation of 0.57 (p-value < 0.001) between the predicted and actual sgRNA growth rates (Figure 3F), suggesting a significant contribution of on-target editing to sgRNA signal. This supports the feasibility of variant validation experiments with pooled sgRNAs by directly sequencing edited variants from genomic DNA.

### Pools of multi-edit sgRNAs create a pool of edited variants that can be measured by error-corrected deep sequencing of genomic DNA

Our BE-HIVE analysis suggested that in-window editing drives multi-edit sgRNA growth rates (Figure 3F). Therefore, we hypothesized that variant validation could be accelerated by directly sequencing variants in genomic DNA during pooled experiments. Instead of validating sgRNA-variant relationships in a one-by-one manner. The proposed approach involves medium-throughput validation pools of sgRNAs, and measuring the resultant variants directly in gDNA via UMI-corrected sequencing of the editing target. To test this, we selected 20 sgRNAs with varying predicted fitness measurements, aiming to determine if a variant is deleterious and to explain sgRNA measurements through direct sequencing of the polyclonal pool (Figure 4A).

**Figure 4.**
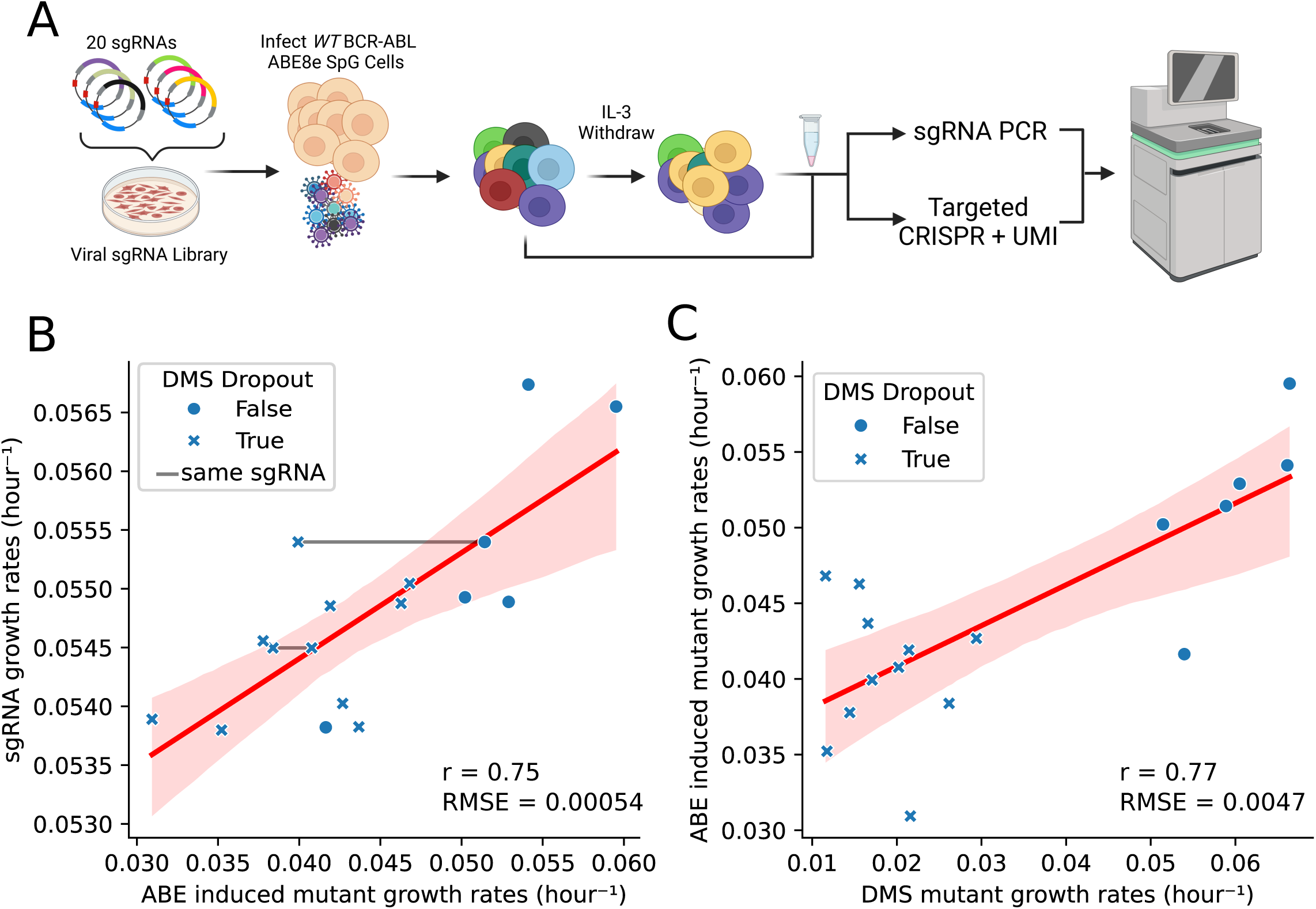
Medium-throughput pooled Adenosine Base Editor Screen. (A) Schematic of small screen of 20 sgRNAs targeting ABL kinase, where the edits and sgRNA are sequenced after IL-3 withdrawal. (B) Comparison of measured sgRNA and single amino acid edit growth rates. If the mutation is deleterious or non-deleterious in the DMS data, then it is marked by a X or closed circle, respectively. If the different mutations are made by the same sgRNA, then they are connected by a grey line. (C) Represents a direct comparison between the measured growth rate of single amino acid mutants during the DMS and 20 sgRNA ABE screens.

The measured sgRNA growth rates strongly correlated (r=0.79, p < 0.001) with the sgRNA growth rates measured in the high-throughput screen from Figure 2, confirming robust phenotypes and validating the new dataset. Further analysis revealed a high correlation (r=0.75) between sgRNA and mutant growth rates (Figure 4B), directly confirming that the fitness effects of sgRNAs are due to on-target editing.

We also examined the correlation between the edited variants growth rate measurements in the medium-throughput pool and the original DMS measurements in Figure 1. Here we found that there was a high correlation (Pearson’s r=0.77) between the direct variant measurements and the gold standard DMS measurements from Figure 1 (Figure 4C). This indicated that the direct sequencing of edited variants in resequencing pools can approach the quality of gold standard DMS data. Among the 19 variants we detected, only F283L showed a spurious depletion phenotype that deviated from observed DMS measurements. This single false positive variant could be due to off-target editing confounding the variant interpretation or be misannotated in the DMS screen. Interestingly, Figure 4C only examined the *single* mutant edits that we detected by deep sequencing, but we also identified double mutants (*in cis*) in multi edit sgRNA windows. Additionally, we identified double mutants in multi-edit sgRNA windows, exhibiting growth rates suggesting no epistasis. The strong correlations and single false positive edit suggest that direct variant sequencing from a base editor pool yields highly concordant fitness measurements with gold standard data (56) (see methods)) (Supplemental Table 2).

## CONCLUSION

In conclusion, our study provides a comprehensive comparison of deep mutational scanning (DMS) and CRISPR-based base editor (BE) screening for variant annotation. We demonstrate that while DMS offers unparalleled depth and structural resolution, BE screening provides a rapid, broad, and efficient alternative at the cost of mutation density. By analyzing both methods in the same cellular and genomic context, we achieved a surprisingly high degree of correlation between the two, despite their inherent methodological differences and the potential for off-target effects in base editing.

Our findings reveal that variant annotation can be achieved directly from base editor screens when focusing on sgRNAs with single predicted edits within a narrow, efficient editing window. This streamlines the annotation process, particularly for variants exhibiting strong phenotypes. We also show that incorporating filters for sgRNA efficiency and reducing multi-edit ambiguity further enhances the correlation between BE and DMS data.

However, for complex cases involving multi-edit sgRNAs and double mutations, direct sequencing of mutant pools offers a robust validation strategy. By directly measuring the variants generated by pooled sgRNAs, we confirmed that the fitness effects observed in BE screens are primarily due to on-target editing. This approach allows for accurate variant annotation even in challenging scenarios, while maintaining a higher throughput.

Overall, our study demonstrates the complementary nature of DMS and BE screening for variant annotation. By strategically combining these two powerful tools, researchers can achieve comprehensive and efficient variant characterization across the genome, accelerating our understanding of gene function and disease mechanisms.

## ACKNOWLEDGEMENTS

We would like to acknowledge members of the Pritchard lab for constructive criticisms of the manuscript, and the Penn State Flow Cytometry Core for help with sorting cells. This work was in part funded by an NSF Modulus Grant MCB 2141650 (J.R.P., D.W.). NIH Grant T32GM108563 (I.S.), NCI U01 synthetic biology grant U01CA265709 (J.R.P.), NSF RECODE grant CBET 2033673 (J.R.P.), NIH grant R35 GM133613 (D.M.M.) and additional funding from the Simons Center for Quantitative Biology at Cold Spring Harbor Laboratory.

---

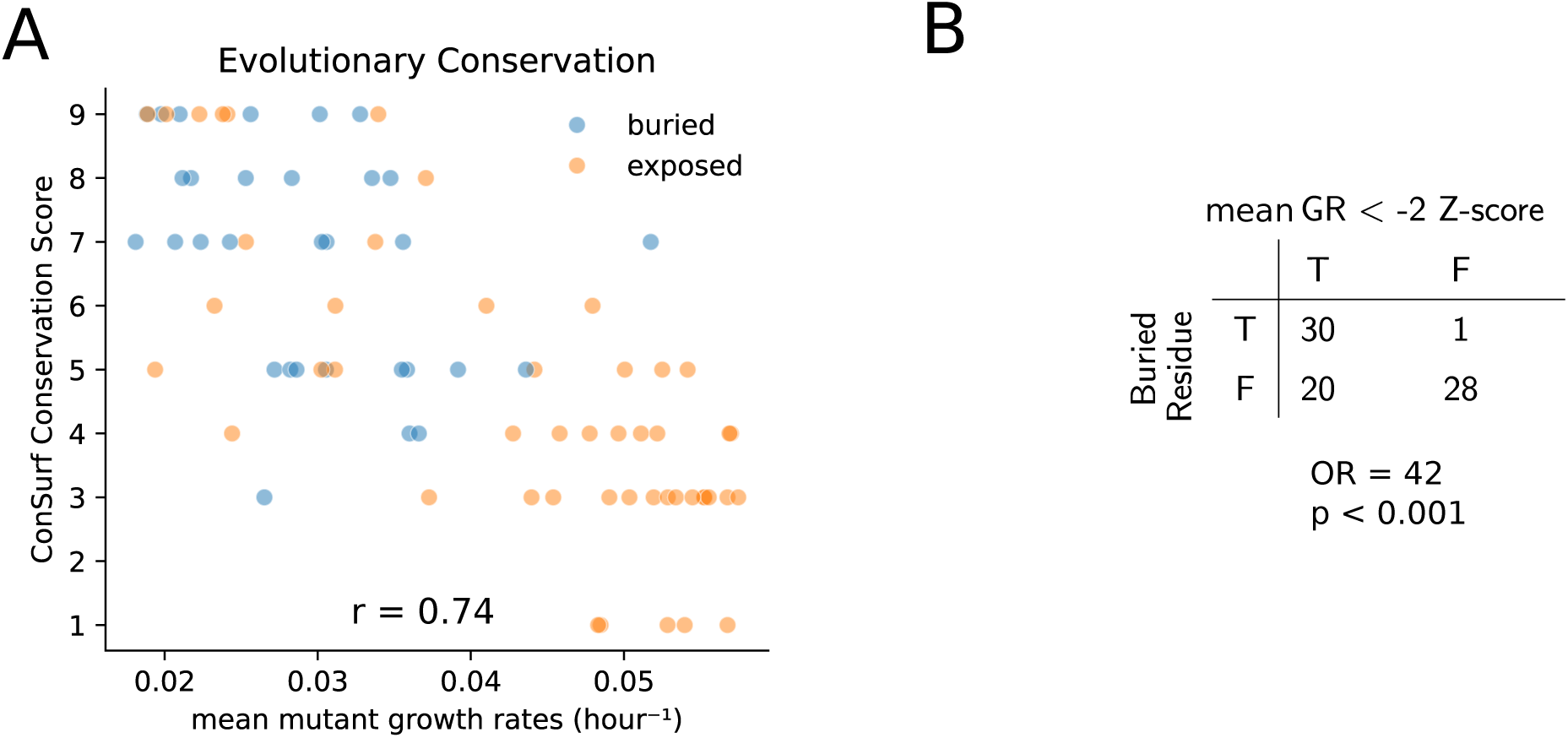

