## Supplemental Figures and Tables for "A side-by-side comparison of variant function measurements using deep mutational scanning and base editing"

**Supplemental Figure 1. Structural and evolutionary conservation is captured by DMS screen.** (A) ConSurf evolutionary conservation score versus mean mutant growth rates for each residue in the ABL N-lobe . Each residue is also color coded based on whether it is exposed (orange) or buried (blue) based on PDB structure 6XR6. (B) Contingency table and fisher exact test statistics comparing residue solvent exposure and if the mean mutant growth rate is less than –2 z-score cutoff drawn from the distribution of WT-like mutant growth rates.

**Supplemental Figure 2. Variant quality control and scoring flow chart.** Excluded variants are boxed in red, and included variants are boxed in blue.

**Supplemental Figure 3. sgRNA quality control and scoring flow chart.** Excluded sgRNAs are boxed in red, and included sgRNA are boxed in blue.

**Supplemental Table 1.** ***Custom Staggered Forward Primers for PCR amplification of sgRNAs from genomic DNA.*** Stagger nucleotides add sequencing diversity. Stagger is bolded and colored purple.

**Supplemental Table 2. Example of multiple edits made by one sgRNA.** Double mutant growth rate is predicted by a product model of mutant interaction.

### Supplemental Figure 1

Sokirniy & Inam

A

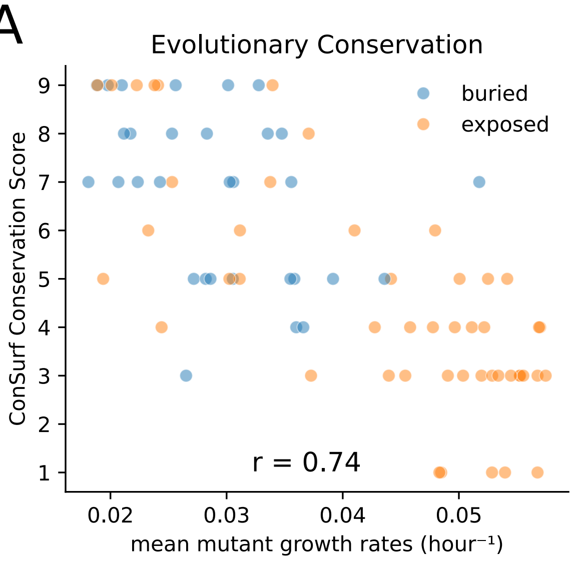


B

| Buried Residue | mean GR < -2 Z-score: T | F |
|---|---|---|
| T | 30 | 1 |
| F | 20 | 28 |

OR = 42
$p < 0.001$

Supplemental Figure 2

Sokirniy & Inam

*If a variant growth rate is less than -2 standard deviations from mean growth rate of the null distribution, then that variant is considered a bona fide hit.*

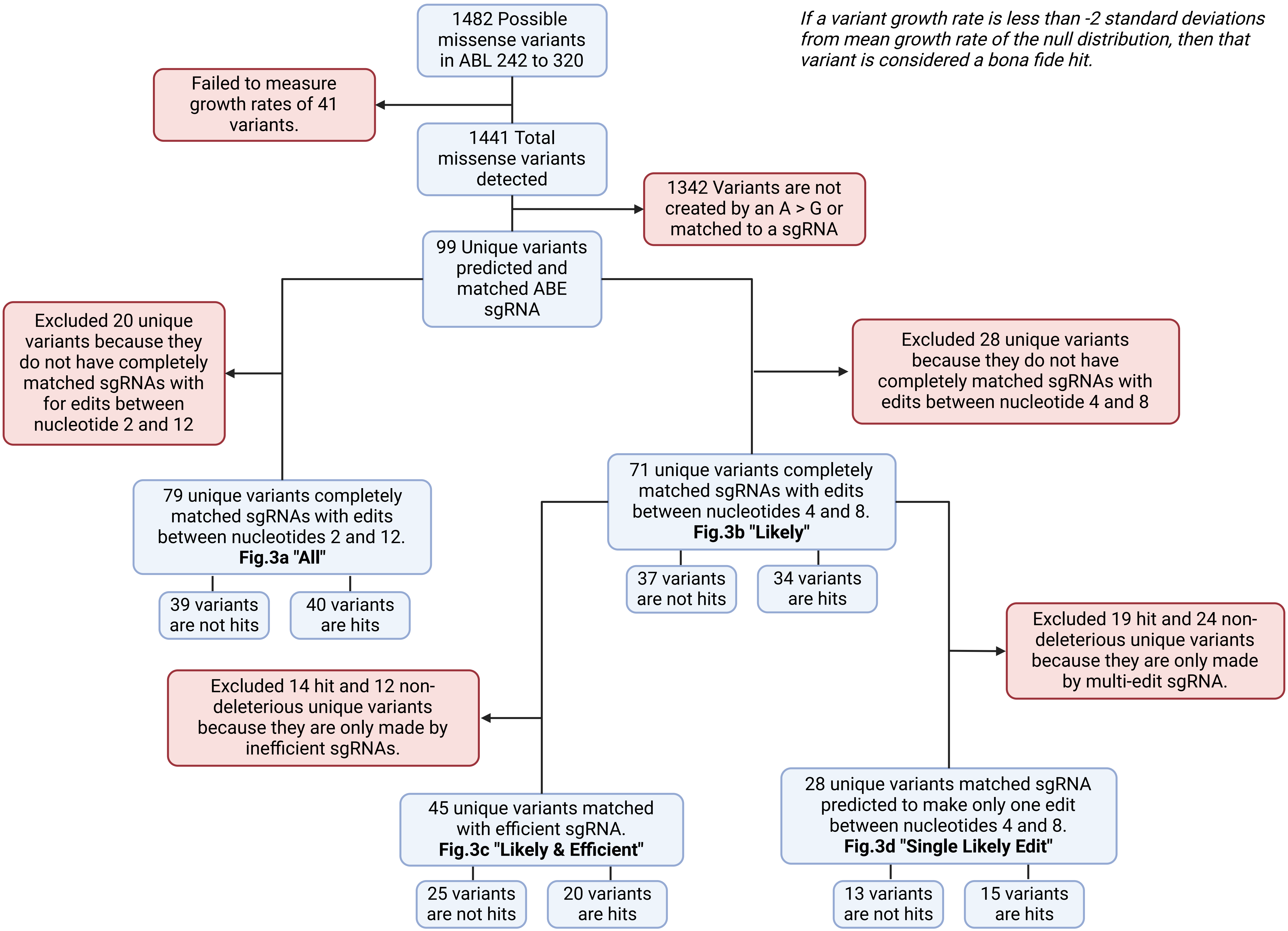

Supplemental Figure 3

Sokirniy & Inam

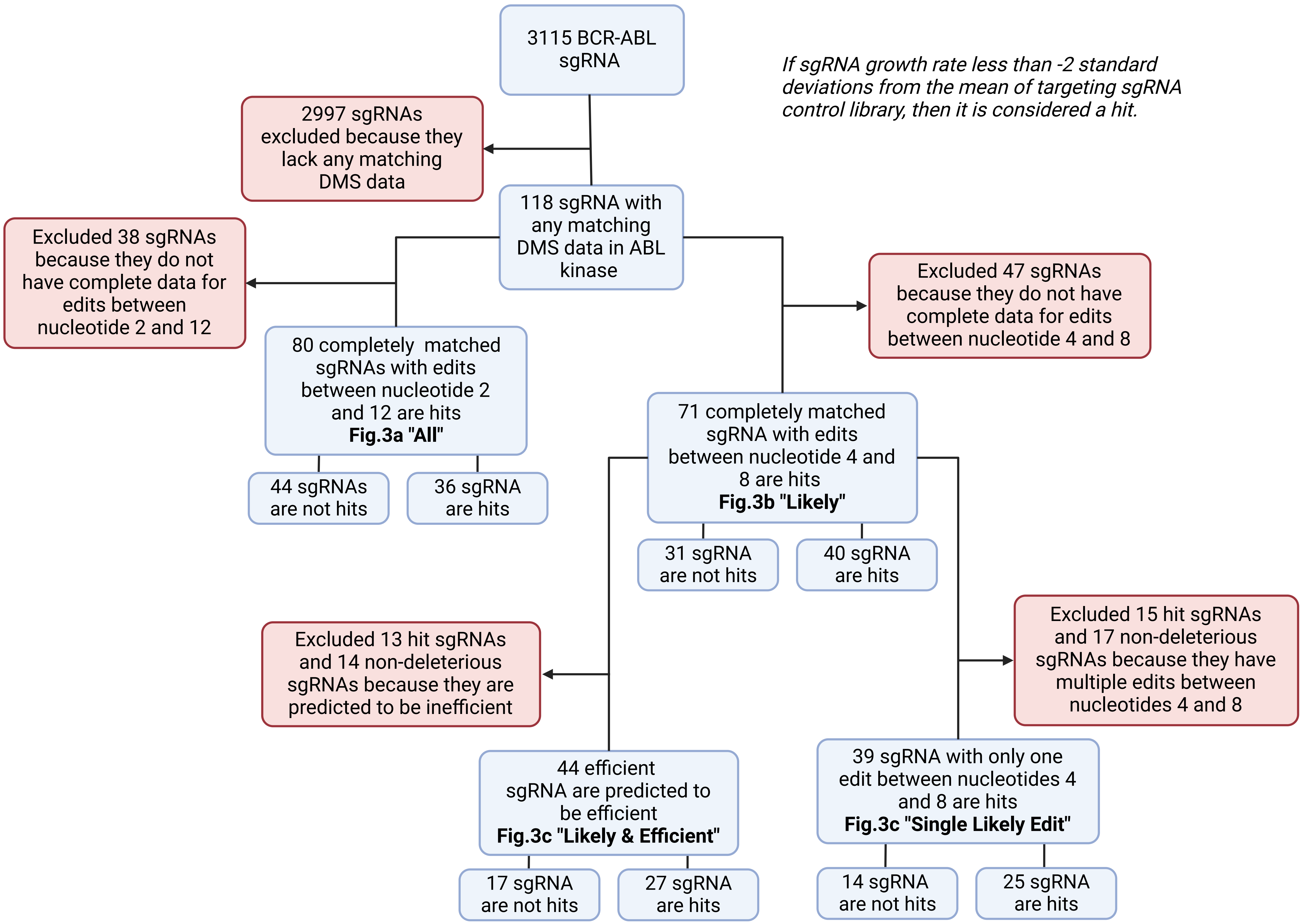

***Custom Staggered Forward Primers for PCR amplification of sgRNAs from genomic DNA***

**Stagger region**

| Primer Name | Sequence |
|---|---|
| NGS_F.n0 | AATGATACGGCGACCACCGAGATCTACACTCTTTCCCTACACGACGCTCTTCCGATC TTGTGGAAAGGACGAAACACC |
| NGS_F.n1 | AATGATACGGCGACCACCGAGATCTACACTCTTTCCCTACACGACGCTCTTCCGATCT **N**TTGTGGAAAGGACGAAACACC |
| NGS_F.n2 | AATGATACGGCGACCACCGAGATCTACACTCTTTCCCTACACGACGCTCTTCCGATCT **NN**TTGTGGAAAGGACGAAACACC |
| NGS_F.n3 | AATGATACGGCGACCACCGAGATCTACACTCTTTCCCTACACGACGCTCTTCCGATCT **NNV**TTGTGGAAAGGACGAAACACC |
| NGS_F.n4 | AATGATACGGCGACCACCGAGATCTACACTCTTTCCCTACACGACGCTCTTCCGATCT **NNVV**TTGTGGAAAGGACGAAACACC |
| NGS_F.n5 | AATGATACGGCGACCACCGAGATCTACACTCTTTCCCTACACGACGCTCTTCCGATCT **NNVVM**TTGTGGAAAGGACGAAACACC |
| NGS_F.n6 | AATGATACGGCGACCACCGAGATCTACACTCTTTCCCTACACGACGCTCTTCCGATCT **NNVVMC**TTGTGGAAAGGACGAAACACC |

**N** = A or C or T or G

**V** = A or C or G

**M** = A or C

***Example of multiple edits made by one sgRNA.***

| Edit | growth rate ($hour^{-1}$) | DMS Pred. growth rate ($hour^{-1}$) |
|---|---|---|
| Q252R | 0.038 | 0.026 |
| Y253C | 0.041 | 0.020 |
| Q252R Y253C | 0.036 | 0.010 |
